# Division of labour underlies efficient colony defence in a clonal ant

**DOI:** 10.1101/2024.02.16.580644

**Authors:** Zimai Li, Qi Wang, Daniel Knebel, Daniel Veit, Yuko Ulrich

**Author notes:** The authors contributed equally to this study (Zimai Li, Qi Wang).

## Abstract

The ecological success of social insects is often credited to the division of labour (DOL), yet empirical evidence directly linking DOL to colony efficiency is rare, in part because variation in DOL between colonies is often confounded by variation in colony traits (e.g., group size, genetic diversity) that can independently affect group efficiency. Here, we measure the link between DOL and efficiency in a crucial task, colony defence, in a social insect that affords precise experimental control over relevant individual and colony traits, the clonal raider ant (*Ooceraea biroi*). We find that DOL in defence behaviour emerges within colonies of near-identical workers, and is consistently associated with enhanced defence efficiency. This positive relationship is robust to variation in the social environment (group size and the presence and type of brood), indicating that the extent of behavioural variation between members of a social group can serve as a key indicator of group efficiency.

## Introduction

Division of labour (DOL), whereby members of a group specialise in distinct tasks, is a key feature of social systems, from microorganisms (West and Cooper 2016) to insects (Wilson 1971, Biedermann and Taborsky 2011) and humans (Smith 1776). DOL plays a key role at all scales of biological organisation (Bonner 1993) and the emergence of new forms of DOL underlies all the major evolutionary transitions, such as those from prokaryotic to eukaryotic cells, from unicellular to multicellular organisms, and from solitary to eusocial life (Queller 1997, Szathmáry and Smith 1995, Bourke 2011). It is widely believed and theoretically predicted that DOL evolved because it increases group efficiency (Beshers and Fewell 2001, Bourke 2011, Duarte et al. 2011, Rueffler et al. 2012, Cooper et al. 2021). However, empirical studies measuring the relationship between DOL and group efficiency remain rare (Oster and Wilson 1978).

Social insects (ants, termites, bees, and wasps) are among the most ecologically successful animals (Wilson 1990, Schultheiss et al. 2022) and display some of the most extreme and elaborate forms of DOL (Wilson 1971, Wilson 1990, Traniello and Rosengaus 1997). Typical social insect colonies have reproductive DOL between one or a few queens that monopolise reproduction and (functionally) sterile workers that perform all other tasks needed for colony maintenance (Hölldobler and Wilson 1990). Additionally, there is (non-reproductive) DOL among the workers, who specialise in a subset of maintenance tasks (e.g., foraging, nursing, defence). Thus, like a multicellular organism, a social insect colony is composed of related units (individuals in insect colonies, cells in multicellular organisms) that are specialised in different essential tasks, like reproduction (queens in social insect colonies, germ line cells in multicellular organisms) or energy storage (repletes in some insect colonies, adipocytes in multicellular organisms), and therefore depend on each other for survival. As a consequence, social insect colonies are often conceptualised as “superorganisms” (Wheeler 1911).

DOL has long been proposed to be the source of the enormous ecological success of ants and other social insects (Oster and Wilson 1978, Wilson 2012) and colonies with high DOL have been hypothesised to be more efficient than colonies with low DOL. A small number of empirical studies have examined the association between DOL and efficiency in e.g., brood rearing (Brahma 2018, Mertl and Traniello 2009, Ulrich 2018), brood rescue (Jongepier and Foitzik 2016), or nest construction (Jeanne 1986), with overall equivocal results. Attempts to quantify the relationship between DOL and efficiency are often complicated by potentially confounding factors such as underlying variation in genetic diversity, age structure, or group size, which can affect both DOL and group efficiency (Robinson 1992, Fewell and Harrison 2016, Ulrich et al. 2018, Brahma 2018, Oldroyd and Fewell 2007, Jeanson and Weidenmüller 2014, Morand-Ferron and Quinn 2011). It thus often remains unclear whether any observed link between DOL and efficiency is driven by DOL itself, or stems from underlying variation in other factors (Jeanson and Weidenmüller 2014).

A striking example of specialisation in social insect workers is in colony defence. Most social insects live in stable nests, which must be defended against various external threats, including conspecific and allospecific intruders and competitors, predators, and parasites (Nouvian and Breed 2021, Barth et al. 2010, Abbot 2022). Analogous to the immune cells that specialise in defending multicellular organisms against external threats like pathogens, individuals in social insects often specialise in colony defence (Cremer and Sixt 2009, Esponda and Gordon 2015).

The best-documented and most iconic cases are the morphologically specialised soldier castes of some termites (Engel et al. 2016), ants (Jaffé et al. 2007), and bees (Grüter et al. 2012). Age polyethism in defence has also been reported, whereby older workers engage in defence more readily than young workers (Oster and Wilson 1978, Robinson 1992, Yanagihara et al. 2018). However, it remains unclear whether DOL in defence can emerge in the absence of inter-individual differences in morphology or age.

Here, we examine the link between DOL and efficiency in colony defence against allospecific intruders in the clonal raider ant *Ooceraea biroi*. In this species, colonies are queenless and composed of workers that reproduce asexually and synchronously, so that genetically near-identical adults are produced in discrete age cohorts (Ravary and Jaisson 2002). This allows us to investigate the relationship between DOL and efficiency in colony defence in the absence of variation in age or genetic background both between individuals and between colonies. We first ask whether there is DOL in colony defence in the clonal raider ant and if so, whether inter-individual variation in defence correlates with variation in exploratory behaviour, as reported in other systems (Bergmüller and Taborsky 2007; Thys et al. 2017, Garamszegi et al. 2009, Sih and Del Giudice 2012, Chapman et al. 2011, Modlmeier et al. 2012, Jandt et al. 2014). We then test whether two aspects of the social environment, colony size and brood type, affect DOL in colony defence. Group size has been shown to increase DOL in other tasks (intranidal vs. extranidal tasks) in the clonal raider ant (Ulrich et al 2018). We expected the presence and type of brood to affect DOL in defence in the clonal raider ant because the brood is known to affect worker physiology and behaviour in this species (Ravary and Jaisson 2002, Fetter-Pruneda et al. 2021). Finally, we ask whether DOL in colony defence correlates with defence efficiency across different experimental conditions.

## Methods

### Experimental design

Clonally related, age-matched (77-day-old) ants of clonal lineage B (Trible et al. 2020) were randomly selected from one stock colony, individually marked on the thorax and gaster using oil-paint markers (Uni Paint PX-20 and PX-21), and used to form experimental colonies with two size treatments (4 workers or 8 workers) and three brood treatments: no brood, larvae (as many 5-day-old larvae as they were workers), or pupae (as many 6-day-old yellow pupae as there were workers). Six replicate colonies were used for each of the six treatments.

Experimental colonies were housed in airtight Petri dishes 5 cm in diameter, with a plaster of Paris floor saturated with water. *O. biroi* feeds on the brood of other ant species (Ravary and Jaisson 2002, Chandra et al. 2021), and colonies were fed *Tetramorium bicarinatum* pupae proportionally to group size (2 pupae for colonies of size 4 and 4 pupae for those of size 8) once. Two days after the colonies were established, we recorded colony behaviour at 20 fps using 6 Basler (model acA20440-20gc; Ahrensburg, Germany) cameras and LoopBio Motif (v6) software. We first recorded one hour of colony baseline activity (Fig. 1A), followed by three trials of colony defence at one-hour intervals. In each trial, we introduced a dead (freeze-killed and thawed) worker of *T. bicarinatum* as an allospecific “intruder” in each colony and recorded a 30-minute video. Pilot experiments were conducted to ensure that the intruder was perceived as an external threat and not as food. Results of the pilots showed that *O. biroi* workers attacked dead workers but not the brood of *T. bicarinatum*, and that dead workers were not consumed, while the brood was. The use of a dead intruder allowed us to measure colony behavioural responses while ruling out any effects of the intruder’s behaviour (Roulston et al. 2003, Modlmeier and Foitzik 2011). Intruders were removed after each trial and a fresh intruder was used in each trial. All experiments were conducted in a climate chamber maintained at 28.20 ± 0.20 °C.

**Figure 1.**
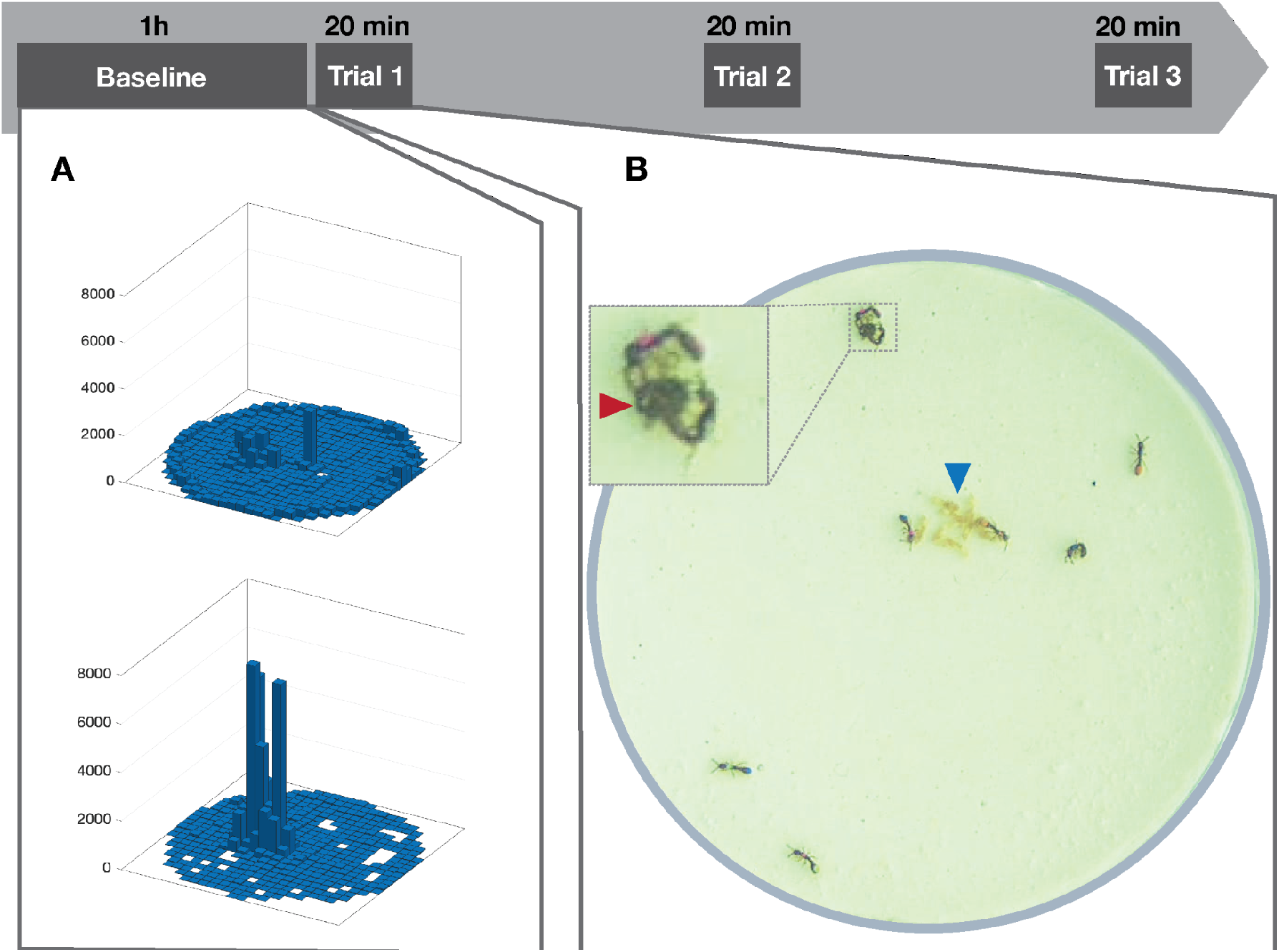
Experimental procedure (A) 3-dimensional histograms of the spatial distribution of one ant with high entropy (top: *H* = 8.415) and one ant from the same colony with lower entropy (bottom: *H* = 3.041) at baseline. The Z-axis represents the number of frames in which the ant occupied each grid cell. (B) Experimental ant colony in a colony defence trial; blue arrow: brood pile (pupae). Insert: zoomed-in view of defence behaviour by two clonal raider ants towards an intruder (red arrow).

### Behavioural data acquisition

Baseline activity in the first hour of recording (before defence trials) was analysed using automated tracking with anTraX (Gal et al. 2020). This detected the position of each ant every 50 ms. Raw tracking data underwent preprocessing in MATLAB (R2023b) to identify missing or aberrant positions, which were then interpolated or removed, respectively, following Jud et al. 2022.

In each trial, we used BORIS (Friard and Gamba 2016) to manually annotate behaviour in the first 20 min after the introduction of the intruder. We annotated encounters of all clonal raider ants (individually identifiable by their unique paint marks) with the intruder as well as their defence behaviour towards the intruder. We quantified the number of encounters, defined as physical contacts with the intruder, as well as the number and duration of stinging attempts (in seconds), defined by a stereotyped holding of the intruder with the mouthparts and bending of the gaster towards the intruder (Fig. 1B; Kronauer et al. 2013).

### Behavioural data analysis

Unless stated otherwise, analyses were conducted using R 4.3.1 (R Core Team 2020). To measure individual baseline exploratory behaviour, we calculated the entropy of individual spatial distribution in the first hour of recording, before defence trials (Hanisch 2023). To this aim, the area of each Petri dish was binned (25 × 25 bins of 2 × 2 mm each; 2 mm is approximately one ant body length). For each ant, we calculated Shannon entropy:

*H*= −Σ *p*(*x*) *log*(*p*(*x*)), where p(x) is the proportion of frames that an ant spent in bin x. Higher entropy corresponds to movement patterns that are more evenly distributed across the grid (indicative of higher exploratory behaviour), while lower entropy values correspond to movement patterns that are more restricted to certain areas of the grid (indicative of lower exploratory behaviour) (Fig. 1A).

To measure an individual’s propensity to engage in defence, for each ant, we calculated a defence score as:

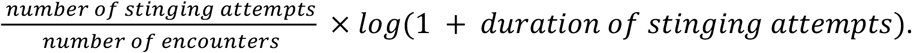

The score incorporates both the number and duration of an individual’s stinging attempts and uses the logarithm of stinging attempt duration to prevent long stinging attempts from inflating the defence score (Roulston et al. 2003). The number of stinging attempts is normalised by the number of encounters to capture the individual propensity to engage in defence behaviour upon encountering an intruder, i.e. to correct for variation in encounter rates (that might arise from, e.g., variation in locomotor activity). The defence score takes the value 0 for ants that never stung the intruder and was assigned the value 0 for ants that never encountered the intruder (Suarez et al. 1999). The defence score correlates with the simpler normalised number of stinging attempts (number of stinging attempts/number of encounters) (Supplementary Figure 1), and statistical analyses using either metric yielded qualitatively similar results (Supplementary Tables 1-4). We calculated the individual defence score of each ant within each trial as well as across all three trials.

To examine the relationship between exploratory and defence behaviour, we tested whether: 1) exploratory behaviour affected encounters with the intruder—as would be expected if ants randomly encountered the intruder while patrolling their environment—using a Gaussian generalised linear mixed model (GLMM; function *glmmTMB* from the *glmmTMB* package), with the number of encounters in trial 1 (a continuous variable) as response variable, individual entropy (a continuous variable), brood treatment (a three-level factor), and colony size (a two-level factor) as predictors, and colony as a random effect; 2) exploratory behaviour correlated with defence scores using as second Gaussian GLMM with the same variables as predictors and individual defence score as response variable (i.e. accounting for inter-individual variation in encounters). We validated the assumptions of both GLMMs using the *simulateResiduals* function from the *DHARMa* package.

To measure DOL in colony defence, we quantified within-individual consistency and between-individual variation in defence behaviour as in Ulrich et al. 2018. Individual consistency in defence behaviour between trials was calculated by ranking individual defence scores within each colony in each trial (assigning the same lowest rank to ties) and conducting Spearman’s rank correlation tests between successive trials (Trial 1 vs. Trial 2 and Trial 2 vs. Trial 3) for each group size and brood treatment. Colonies that did not show defence behaviour in either trial of each pair were excluded from the analysis. To quantify within-colony variation in defence behaviour, we calculated the standard deviation of individual defence scores for all ants in a colony, in each trial separately as well as across all three trials. To investigate factors influencing behavioural variation in colony defence, we used a Gaussian GLMM (*lmer* function from package *lme4*) to analyse the effects of brood treatment (a three-level factor), group size (a two-level factor), and trial (a three-level factor), as well as all their interactions, on behavioural variation, with colony as a random effect. We used the *drop1* function (package *stats*) to evaluate the significance of terms and to reduce models by iteratively deleting non-significant interactions. We validated model assumptions using the *simulateResiduals* function from the *DHARMa* package. Additionally, to assess the impact of group size on behavioural variation while avoiding artefacts arising from sampling effects we employed a resampling approach following Ulrich et al. 2018. We simulated colonies of size 4 by randomly selecting 4 individuals (without replacement) from each colony of size 8. Behavioural variation was calculated for each simulated colony and averaged across replicate simulated colonies. This resampling procedure was repeated 1,000 times. To evaluate whether the behaviour variation of colonies of size 4 significantly differed from that of size 8, we generated 95% confidence intervals for the behavioural variation of simulated colonies using the resampled data.

For each colony, defence efficiency was defined as the total number of stinging attempts received by the intruder across all trials. This is based on the assumption that a higher number of stings is more likely to lead to the retreat or death of intruders and therefore represents a more efficient defence (Gu et al. 2021). We assessed the effects of within-colony inter-individual variation in defence behaviour in all trials (a continuous variable), group size (a two-level factor) and brood treatment (a three-level factor), as well as all their interactions, on colony defence efficiency using a linear regression model (LM, *lm* function from package *stats*). We used the *drop1* function to evaluate the significance of terms and to reduce models by iteratively deleting non-significant interactions. We then performed pairwise comparisons between the levels of significant factors, using Tukey’s posthoc tests (function *emmeans* from package *emmeans*). Model assumptions of the LM were validated with the diagnostic plots produced by the *autoplot* function from package *ggfortify*.

## Results

Individual baseline exploratory behaviour (Figure 1A) was positively associated with the number of encounters with the intruder in the first defence trial (GLMM, DF = 1, LRT = 11.279, p = 0.0008), as expected if clonal raider ants randomly encountered the intruder while patrolling their environment. However, there was no correlation between individual baseline exploratory behaviour and individual defence score (GLMM, LRT = 2.23, p = 0.1357). Thus, while exploratory behaviour increased the likelihood of encountering intruders, it did not increase the propensity of an ant to engage in defence upon encounters.

We find evidence for DOL in defence behaviour. Within a colony, individual defence scores could range from 0 (in an ant that encountered the intruder 14 times and never stung) to 4.66 (in an ant that stung the intruder in 13 out of 18 encounters for a total duration of 629.25 seconds) in one trial. Individual defence scores were overall positively correlated across trials, demonstrating individual consistency in defence behaviour (Fig. 2A, Supplementary Figure 2, Supplementary Table 5; group size 4: Trial 1 vs. Trial 2: R = 0.17, p = 0.1950, Trial 2 vs. Trial 3: R = 0.41, p = 0.0040; group size 8: Trial 1 vs. Trial 2: R = 0.28, p = 0.0008; Trial 2 vs. Trial 3: R = 0.46, p = 5.52 × 10^−9^). DOL in defence increased with group size. Individual consistency in defence behaviour was overall stronger in larger colonies (see above; Fig. 2A). Similarly, variation in defence behaviour was higher in larger colonies (Fig. 2C, GLMM: DF = 1, sum Sq = 1.79, F = 9.34, p = 0.0045) and this effect could not be explained by sampling effects, as shown by the fact that the observed values for small colonies (mean ± s.e.m in colonies of size 4: 0.822 ± 0.080) fell outside of the confidence interval generated by resampling from large colonies (95 % confidence interval in simulated small colonies resampled from large colonies: (0.923, 1.170)). Finally, DOL (behavioural variation) decreased across trials (DF = 2, Sum sq = 3.01, F = 7.88, p = 0.0008; Supplementary Table 6) but was not affected by brood treatment (DF = 2, sum Sq = 0.409, F = 1.07, p = 0.3548).

**Figure 2.**
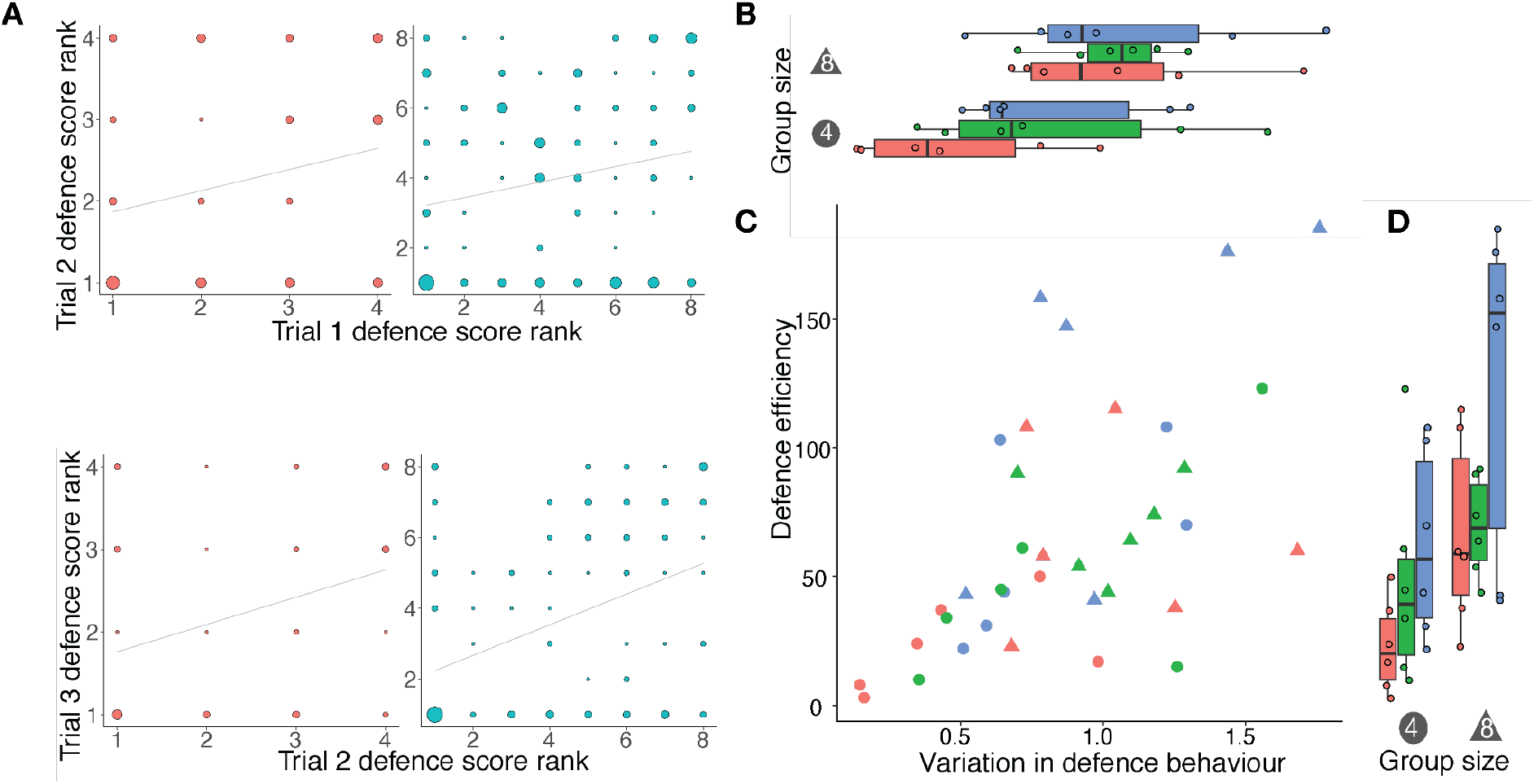
Division of labour in colony defence. (A) Behavioural consistency: correlation between ranked defence scores in trial 1 vs. trial 2 (top) and trial 2 vs. trial 3 (bottom). Data points represent individuals; data point diameter is proportional to the number of ants with a given rank combination; colours represent group size (red: 4, blue: 8); grey lines: least-squares fit. Data for different brood treatments are pooled. (B) Variation in defence behaviour in all trials as a function of group size and brood treatment (red: no brood, green: larvae, blue: pupae). Data points represent colonies; boxes indicate the interquartile range (IQR) around the median (thick black line), whiskers represent the highest and lowest values within 1.5 * IQR of the upper and lower quartiles, respectively). (C) Colony defence efficiency as a function of behavioural variation. Colours as in B; symbols represent group size (circles: 4, triangles: 8) (D) Defence efficiency across all trials as a function of group size and brood treatment (colours as in B). Data points represent colonies; boxes indicate the IQR around the median (thick black line), whiskers represent the highest and lowest values within 1.5 * IQR of the upper and lower quartiles, respectively.

Colony defence efficiency increased with within-colony variation in defence behaviour (Fig. 2C, LM, DF = 1, Sum Sq = 10261.30, F = 8.08, p = 0.0078; Supplementary Figure 3) robustly across group sizes (interaction between variation in defence behaviour and group size: DF = 1, Sum Sq = 5.70, F = 0.0044, p = 0.9478) and brood treatments (interaction between variation in defence behaviour and brood treatments: DF = 2, Sum Sq = 5380.20, F = 2.30, p = 0.1187). Thus, there was a consistent positive relationship between DOL in defence and defence efficiency across conditions. Additionally, colony defence efficiency was influenced by the presence and type of brood (Figure 2D, DF = 2, Sum Sq = 4.40, F = 4.02, p = 0.0207). Colonies with pupae had higher defence efficiency than those without brood (Tukey’s posthoc test: no brood - pupae, estimate= -40.14, SE = 14.90, df = 31, t =-2.70, p = 0.0292), while colonies with larvae did not differ from colonies without brood (no brood - larvae, estimate = -5.24, SE = 14.90, df = 31, t = -0.35, p = 0.9339) or from colonies with pupae (larvae - pupae, estimate = - 34.90, SE = 14.50, df =31, t = -2.40, p = 0.0570). Finally, colony defence efficiency was only marginally affected by group size (DF = 1, Sum Sq = 5274.70, F = 4.15, p = 0.0501).

## Discussion

We found DOL in colony defence in *O. biroi*, as demonstrated by inter-individual variation and individual consistency in defence behaviour. Although all ants used here had near-identical age and genotype and were reared in the same controlled environment, they differed consistently in defence behaviour, suggesting that DOL in colony defence can arise in small, homogeneous social groups. Because we measured the performance of a task in response to a controlled experimental stimulus and corrected for variation in encounter rates with the stimulus, the observed differences in behaviour are likely to reflect individual variation in response thresholds. Variation in response thresholds is one of the theoretically best-studied mechanisms by which DOL can emerge (Beshers and Fewell 2001, Bonabeau and Theraulaz 1999) but empirically measured variation in response thresholds is still rare. Here, we detect consistent variation in response thresholds that is independent of age or genetic background (as shown by inter-trial consistency in defence scores). As shown in the same species for other behaviours (Ulrich et al., 2018), DOL in defence increased with colony size. Additionally, colonies with higher variation in defence behaviour also had more efficient defence (i.e., they collectively inflicted a higher number of stings on the intruder). Although this association remains correlational, the correlation was consistent across experimental conditions (group size, presence and type of brood), suggesting a direct link between DOL and efficiency, instead of one driven by co-varying and thus potentially confounding factors such as variation in demographic/genetic diversity or group size. The positive link between DOL and efficiency detected here mirrors similar findings in other systems, including cooperatively breeding birds (Ridley and Raihani 2008), other social insects (Mertl and Traniello 2009, Brahma 2018, Jeanne 1986), and even parasitic trematodes (Lloyd and Poulin 2012). However, the link may not be universal. For example, experimental increases in specialisation in colonies of acorn ants negatively affected brood rescue efficiency during attacks (Jongepier and Foitzik 2016). This suggests that while the relationship between DOL and group efficiency may overall be positive, above certain levels, or under certain conditions, specialisation can also hinder a group’s flexibility and decrease its efficiency, as predicted theoretically (Cooper and West 2018).

We failed to detect a behavioural syndrome linking exploration and aggression as joint manifestations of boldness that correlate across individuals, as has been reported in various species (Bergmüller and Taborsky 2007; Thys et al. 2017, Garamszegi et al. 2009, Sih and Del Giudice 2012), including ants (Chapman et al. 2011, Modlmeier et al. 2012, Jandt et al. 2014). Here, while ants with higher explorative behaviour encountered intruders more frequently, they were not more (or less) likely to engage in defence behaviour upon encounter. Past work on social insects involved colonies containing individuals of varying ages, genetic backgrounds, and/or morphology. In contrast, we controlled for age and genetic background in *O. biroi*, a monomorphic species, and found no correlation between exploratory and defensive behaviours. This suggests that the behavioural syndrome reported in social insects might stem from underlying inter-individual variation in factors such as genotype (Alaux et al. 2009), caste (Chapman et al. 2011), or age (Judd 2000).

The type of brood present in colonies affected defence efficiency, with colonies containing pupae displaying more efficient defence than colonies without brood. This pattern could reflect an “economic strategy” of the colony (Cole 1988, Sakata and Katayama 2001). Pupae used in this study were the product of ca. 20 days of brood care, a substantial time and resource investment. In addition, late-stage ant pupae have nutritional value for the colony: the moulting fluid they produce is rich in nutrients, hormones, and neuroactive substances that are consumed by adults (Snir et al. 2022). It is therefore plausible that the high value of pupae triggers a more efficient defence response than in colonies without brood.

Our results show that colony efficiency hinges on variation in individual behavioural responses. Similar phenomena exist in many social groups, including thermoregulation in bees (Jones et al. 2006, Weidenmüller 2004), cooperative transport in ants (Feinerman et al. 2018), group foraging in guppies (Dyer et al. 2009), collective cell movement (Blanchard 2019), and information spread in humans (Zhu et al. 2014). While the magnitude of a response may not always reflect its efficiency at the group level (excessive behavioural responses can also lead to catastrophic failure in e.g., stampedes (Helbing et al. 2007)), for tasks such as defence, the group-level magnitude of the behavioural response (e.g. stinging attempts) is likely to be a reasonable proxy for its efficiency (e.g. the likelihood to successfully repel or kill an intruder) (Gu et al. 2021). Our findings indicate that in such scenarios, the extent of behavioural variation between members of a social group can serve as a key indicator for the efficiency of the resulting global response.

## Data accessibility

The data and code for analysis are available from GitHub repository: https://github.com/lizimai/Li_etal_2024.

## Author’s contributions

Z.L.: conceptualisation, data acquisition, data curation, methodology, supervision, investigation, writing - original draft and writing - review and editing; Q.W.: conceptualisation, data acquisition, methodology, investigation; D.K.: conceptualisation, methodology, supervision, writing - review and editing; D.V., experiment setup; Y.U.: funding acquisition, project administration, methodology, supervision, validation, writing - review and editing.

All authors gave final approval for publication and agreed to be held accountable for the work performed therein.

## Conflict of interest declaration

We declare we have no competing interests.

## Acknowledgement

The authors thank Baptiste Piqueret for suggestions on experimental design, and Lai Ka Lo, Bhoomika Bhat, Sandra Tretter, Xiaohua Chu, Sarah Rogoz and Tim Zetzsche for discussions on the results. This work was supported by the Max Planck Society. We thank Michael and Barbara Taborsky for organising the workshop “Division of labour as key driver of social evolution” at the Wissenschaftskolleg zu Berlin in March 2023 and for inviting us to contribute to this special issue. This is Clonal Raider Ant Project paper #33.

---

**Supplementary Table 1.** Spearman correlation tests on the individual rank of normalised stinging attempts (number of stinging attempts/number of encounters) between trials for different colony sizes.

| **Trial comparison** | **Group Size** | **r** | **p** |
| --- | --- | --- | --- |
| Trial 1 vs 2 | 4 | 0.23 | 0.07584141 |
| Trial 2 vs 3 | 4 | 0.33 | 0.02280048 |
| Trial 1 vs 2 | 8 | 0.20 | 0.01451963 |
| Trial 2 vs 3 | 8 | 0.42 | 1.42E-07 |

**Supplementary Table 2.** Spearman’s correlation test on ranked individual normalised stinging attempts (number of stinging attempts/number of encounters) between successive trials for brood treatments.

| **Trial comparison** | **Brood condition** | **r** | **p** |
| --- | --- | --- | --- |
| Trial 1 vs 2 | No brood | 0.31 | 0.0144164 |
| Trial 2 vs 3 | No brood | 0.45 | 0.00027908 |
| Trial 1 vs 2 | Larvae | 0.20 | 0.10018932 |
| Trial 2 vs 3 | Larvae | 0.51 | 4.46E-06 |
| Trial 1 vs 2 | Pupae | 0.33 | 0.00436489 |
| Trial 2 vs 3 | Pupae | 0.26 | 0.04587796 |

**Supplementary Table 3.** Effect of group size, brood treatment, and trial on normalised stinging attempts (number of stinging attempts/number of encounters)

|  | **Sum Sq** | **DF** | **F** | **p** |
| --- | --- | --- | --- | --- |
| group size | 0.074316 | 1 | 8.7113 | 0.005877 |
| brood | 0.02461 | 2 | 1.4424 | 0.251313 |
| trial | 0.012252 | 2 | 0.7181 | 0.491244 |

**Supplementary Table 4.**
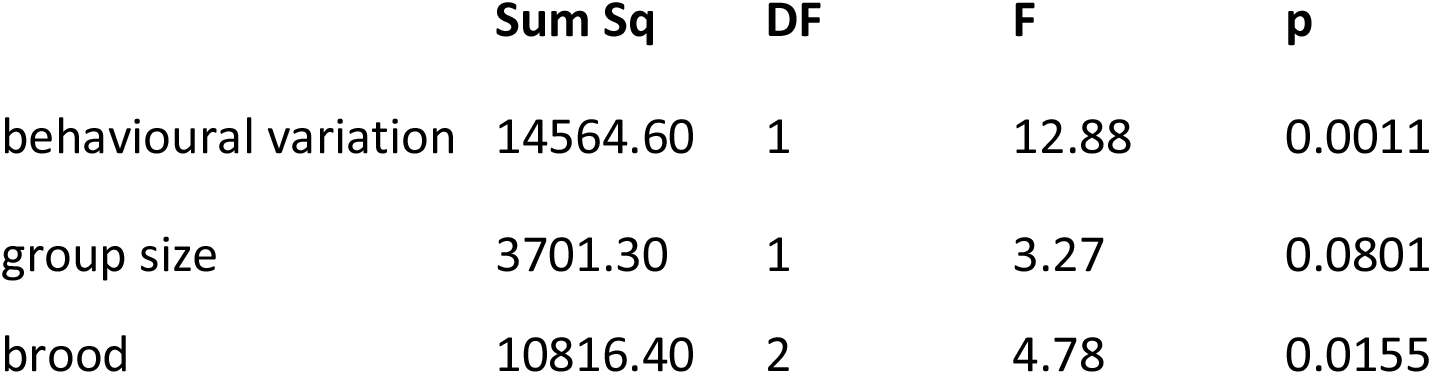
Effect of variation in normalised stinging attempts, group size, and brood treatment on defence efficiency.

|  | **Sum Sq** | **DF** | **F** | **p** |
| --- | --- | --- | --- | --- |
| behavioural variation | 14564.60 | 1 | 12.88 | 0.0011 |
| group size | 3701.30 | 1 | 3.27 | 0.0801 |
| brood | 10816.40 | 2 | 4.78 | 0.0155 |

**Supplementary Table 5.**
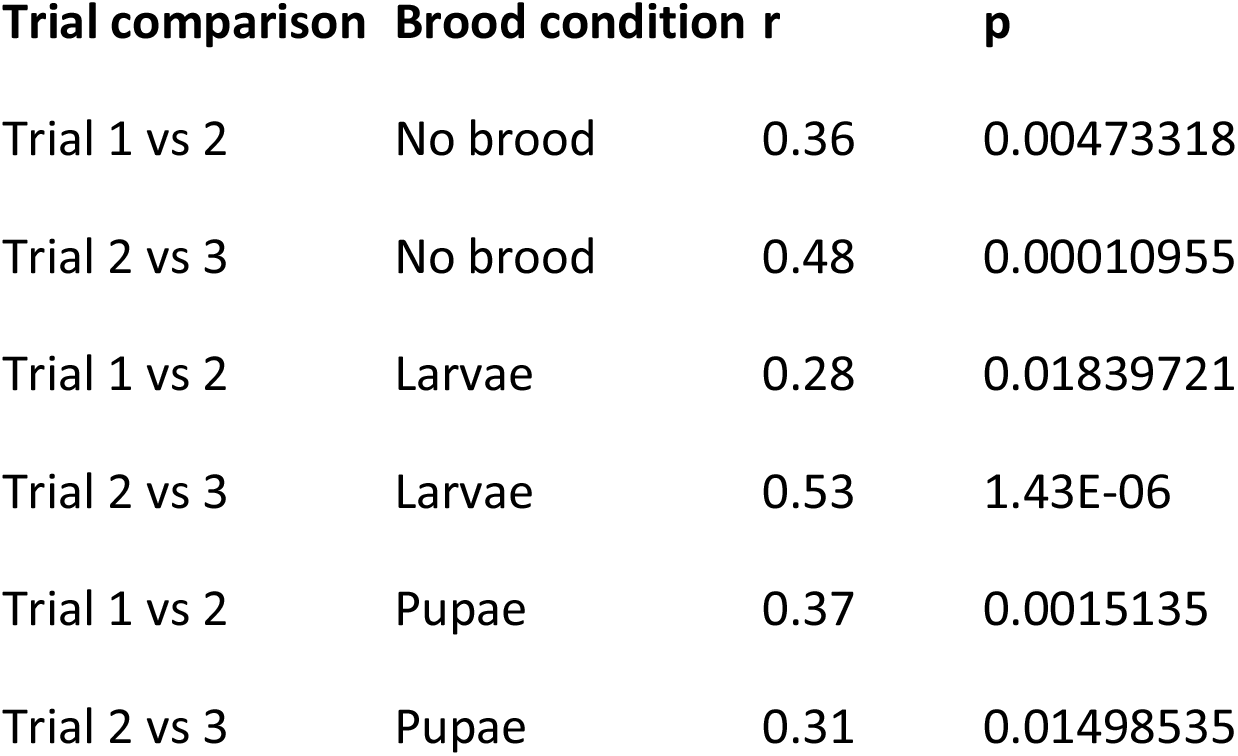
Spearman’s correlation tests on ranked individual defence scores between successive trials for different brood treatments.

| **Trial comparison** | **Brood condition** | **r** | **p** |
| --- | --- | --- | --- |
| Trial 1 vs 2 | No brood | 0.36 | 0.00473318 |
| Trial 2 vs 3 | No brood | 0.48 | 0.00010955 |
| Trial 1 vs 2 | Larvae | 0.28 | 0.01839721 |
| Trial 2 vs 3 | Larvae | 0.53 | 1.43E-06 |
| Trial 1 vs 2 | Pupae | 0.37 | 0.0015135 |
| Trial 2 vs 3 | Pupae | 0.31 | 0.01498535 |

**Supplementary Table 6.**
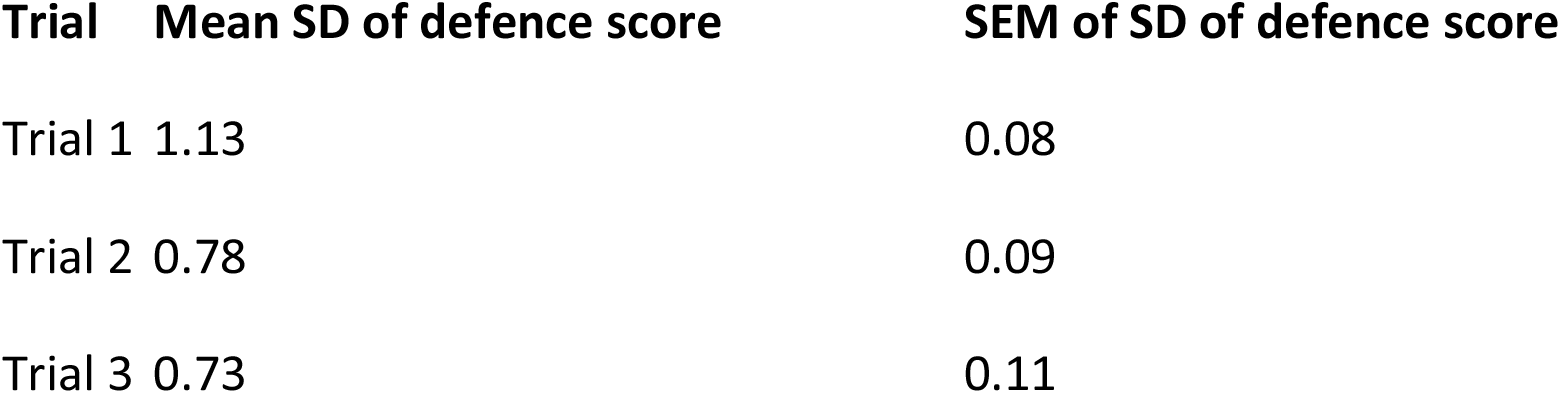
Behaviour variation of defence scores in trial 1, trial 2, and trial 3, colonies from different brood treatments and group sizes are pooled.

**Supplementary Figure 1.**
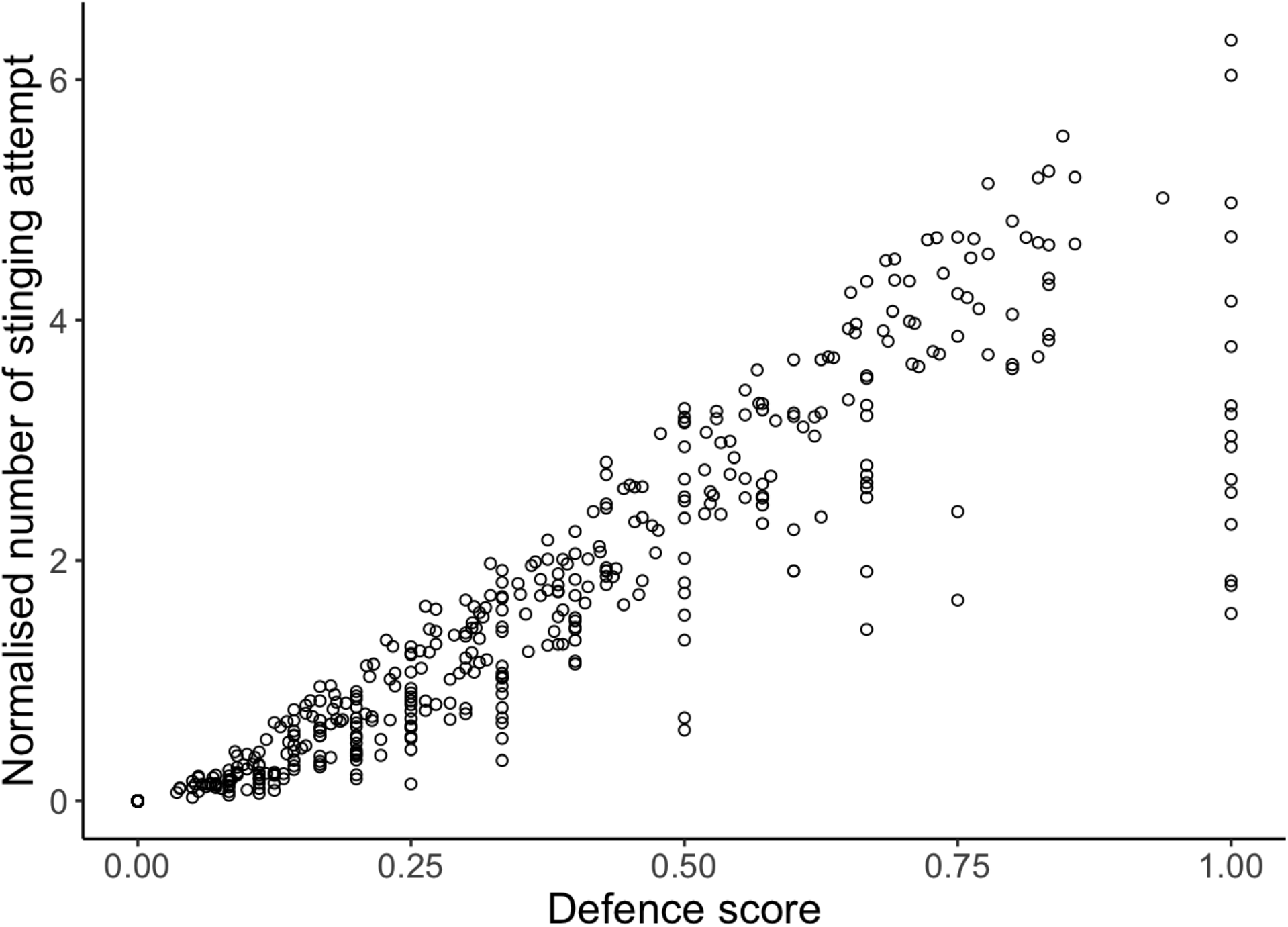
Correlation between individual defence score and normalised stinging attempts (number of stinging attempts/number of encounters). Circles represent individuals in each of the three trials (Pearson correlation test: t = 67.50, df = 646, p < 0.0001, r = 0.94).

**Supplementary Figure 2.**
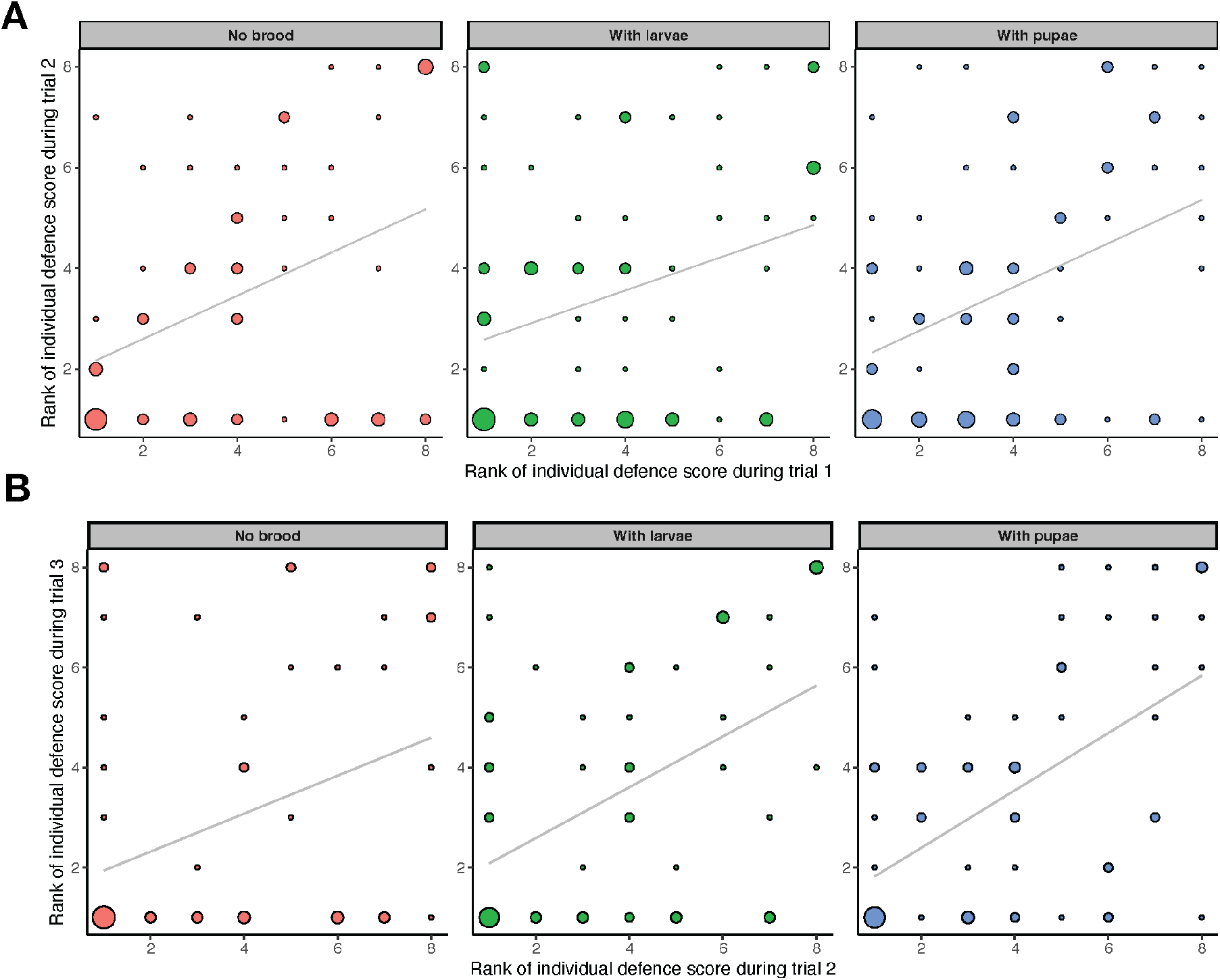
Correlation between individual ranks of defence scores in trial 1 and trial 2 and trial 2 and trial 3. The circle diameter is proportional to the number of ants for a given rank combination; grey lines: least-squares fit. Data for different group sizes are pooled.

**Supplementary Figure 3.**
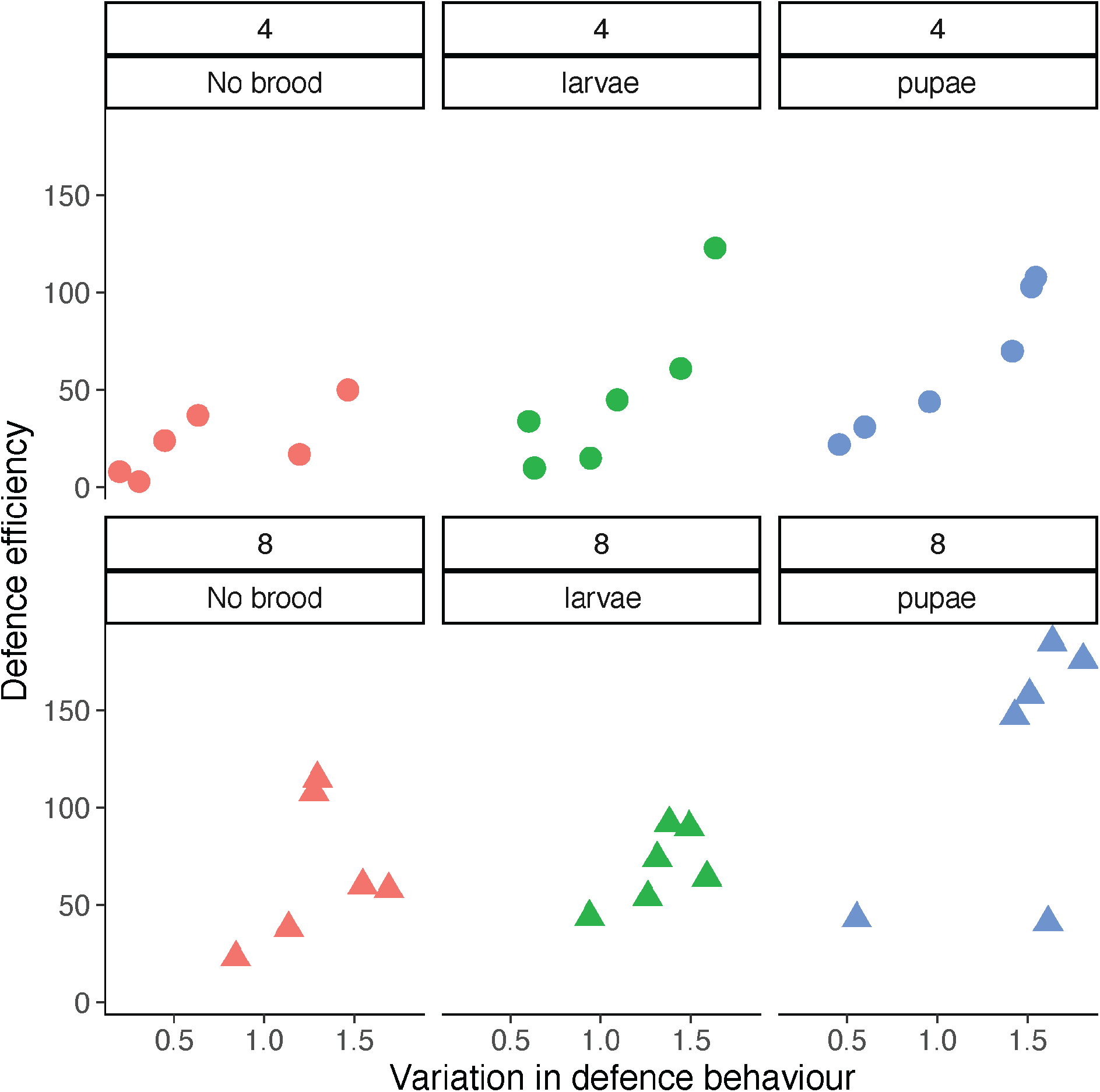
Defence efficiency as a function of the behavioural variation, colours represent brood condition (red: no brood, green: larvae, blue: pupae) and shapes of the dots represent group size (circle: 4, triangle: 8).

## Notes

### Competing Interest Statement

The authors have declared no competing interest.

